# Making new connections: An fNIRS machine learning classification study of neural synchrony in the default mode network

**DOI:** 10.1101/2025.05.31.656874

**Authors:** Grace Qiyuan Miao, Ian J. Lieberman, Ashley L. Binnquist, Agnieszka Pluta, Bear M. Goldstein, Rick Dale, Matthew D. Lieberman

## Abstract

Successfully making connections with others is crucial to navigating the social world and general well-being, yet little is known about connection formation and its neurocognitive underpinnings. Increasingly, neuroscientists use interpersonal ‘neural synchrony’ within the default mode network (DMN) to measure when two or more people subjectively experience something in similar ways. DMN synchrony as ‘seeing eye-to-eye’ is typically observed when multiple people are passively observing the same stimulus. In this study, we tested whether the same DMN synchrony as ‘seeing eye-to-eye’ pattern holds during social interactions. We conducted a between-subject naturalistic experiment with 70 pairs of strangers engaged in either shallow or deep conversations while brain activity was measured with functional near infrared spectroscopy (fNIRS). Stranger dyads successfully formed connections, as indicated by composite connection scores. Replicating Kardas et al. (2021), those in the deep conversation condition felt more connected than those in the shallow conversation condition. DMN neural synchrony significantly predicted self-reported connection, with synchrony in the DMN subregions of medial prefrontal cortex (mPFC) and right temporoparietal junction (TPJ) each correlating significantly with connection. Using machine learning classification, we distinguished high-versus low-connection dyads based on DMN neural synchrony and the perceived depth of conversation with 64.5% accuracy across 1,000 iterations. This effect was primarily carried by right TPJ, which alone classified connection strength at 62.6% accuracy. We consider implications related to the growing loneliness crisis and the importance of understanding how social connections can be formed and fostered in an era of increased social isolation.

**Significance Statement:** The current loneliness epidemic has serious consequences on health and well-being. Forming interpersonal connections is crucial for alleviating loneliness, yet little is known about its neural basis. Neural synchrony, a potential biological marker of people being on the ‘same page’, may be an indicator of social connection. We recorded brain activity as strangers engaged in a get-to-know-you conversation and found that neural synchrony—specifically within the default mode network (DMN) and subregions including medial prefrontal cortex (mPFC), and right temporoparietal junction (TPJ)—predicted self-reported connection. Machine learning accurately classified high- and low-connection dyads based on DMN synchrony and perceived conversation depth. These findings suggest that deeper conversations, though more effortful, may foster stronger social bonds with measurable neural correlates.

## Introduction

The current loneliness epidemic has major negative consequences for public health and well-being (Murthy, 2020; Park et al., 2020; Hong et al., 2023). Increasing social connections with others is crucial to addressing loneliness (Masi et al., 2011), yet little is known about connection formation and its neurocognitive bases.

Increasingly, neuroscientists use interpersonal ‘neural synchrony’ within the default mode network (DMN) as a measure of when two or more people subjectively experience something in similar ways (i.e., ‘seeing eye-to-eye’; Lieberman, 2022, in press). The decision to focus on the DMN in this context is motivated by its hypothesized role as a “sense-making” network, integrating extrinsic information with prior knowledge to construct rich, context-dependent internal narratives (Yeshurun, Nguyen, & Hasson, 2021; Lieberman, 2022; Meyer, 2019; Mars et al., 2012; Menon, 2023).

Investigating DMN activity when individuals watch naturalistic stimuli (e.g. movie clips), multiple functional magnetic resonance imaging (fMRI) studies have demonstrated that greater neural synchrony within DMN regions is linked to both shared ways of experiencing (Yeshurun et al., 2017; Saalasti, 2019; Nguyen, 2019; Dieffenbach et al., 2021) and stronger real-world social connections including friendship and marriage (Parkinson, Kleinbaum, & Wheatley, 2018; Li et al., 2022). Additionally, a recent study by Speer et al. (2024) examined brain responses between friends and stranger dyads during conversations, finding that friends’ brain activity diverged in ‘neural state space’, while strangers’ brain activity converged. Their study revealed the potential role of convergence in dyadic interaction. However, some of the observed differences between friends and strangers likely reflected pre-existing connections between friends, rather than changes in closeness generated by the conversation itself. Conversation has been studied more extensively with functional near infrared spectroscopy (fNIRS) (Kelsen et al., 2022). However, none of these fNIRS studies have examined feelings of connection engendered by a conversation.

Our goal was to examine whether DMN synchrony during active conversations predicts social connection, similar to findings in passive observation studies (Parkinson, Kleinbaum, & Wheatley, 2018; Li et al., 2022). If DMN synchrony reflects two people seeing eye-to-eye as they talk, this ought to predict self-reported feelings of connection afterwards. To examine this, we ask strangers to come into the lab and engage in a get-to-know you conversation while fNIRS was used to record their neural activity. To encourage more variability in how connected participants felt after the conversation, dyads were assigned either shallow or deep prompts to guide their conversations, as prior literature suggests that deep questions tend to induce greater connection (Kardas, Kumar, & Epley, 2021). We hypothesized that DMN synchrony and how deep the conversation felt would predict the degree of self-reported connection present between strangers. We tested this both by correlating DMN neural synchrony and felt depth with self-reported feelings of connection after the conversation and also by conducting machine learning-based classification analyses. As our fNIRS montage also captured data from other neural networks including the frontoparietal network (FPN), the dorsal attention network (DAN), and the ventral attention network (VAN), we examined each of these as well, but had no *a priori* hypotheses for these networks.

## Materials and Methods

### Participants

We ran our procedure on 210 individuals who formed 105 stranger dyads (23 male-male, 38 female-female, 39 male-female, 3 female-other, and 2 male-other). Behavioral analysis is based on all 105 dyads, while neuroimaging results are based on a sample of 70 dyads (20 male-male, 24 female-female, 23 male-female, 2 male-other, and 1 female-other) due to technical issues with the hardware that was identified after the collection of the first 35 dyads. Behavioral analyses using the 70 dyads from the fNIRS analyses show the same patterns as the 105 dyads, but with less statistical power.

Participants were recruited from the UCLA Departments of Psychology and Communication subject pools as well as flyers on the UCLA campus. Experimental procedures were approved by the UCLA Institutional Review Boards (#22-001209) and participants provided informed consent prior to the start of the study. Participants received either 2 course credits or a $30 gift card for the 2-hour long experiment. All participants self-reported as having English as their primary language.

### Experimental procedures

This was a between-subject experimental design with two conditions: shallow and deep (among 105 dyads for behavioral analysis – shallow condition: 12 male-male, 20 female-female, 19 male-female, and 2 male-other; deep condition: 11 male-male, 18 female-female, 20 male-female, and 1 female-other; among 70 dyads for neuroimaging analysis – shallow condition: 10 male-male, 13 female-female, 11 male-female, and 2 male-other; deep condition: 10 male-male, 11 female-female, 12 male-female, and 1 female-other).

In each session, two strangers engaged in a face-to-face, get-to-know-you conversation, with either shallow or deep topics displayed sequentially on a computer screen (Fig 1 in Miao et al. (2023)). During the experiment, participants wore a functional near-infrared spectroscopy (fNIRS) rig. Three GoPro cameras were placed in the room to record conversations and nonverbal behaviors. One camera was positioned in front of each participant to capture facial expressions, and a third camera captured both participants together. Participants also completed questionnaires regarding their personal traits and experiences with the conversation.

The conversation topics (Table 1) were adapted from Kardas, Kumar, & Epley (2022). These topics appear one-by-one in the same order on a computer screen accessible to both participants. Participants read the question at the same time and volunteered to answer in the order they preferred. Participants were instructed to stay on topic, take however much time they needed for every prompt, and when they finished discussing one topic, click a button to move on to the next. Each session was designed to last 20 minutes and took place without the presence of experimenters, allowing for a more natural conversation flow.

**Table 1.** Conversation topics assigned to participants.

| Conversation Topics |  |
| --- | --- |
| Shallow Condition | 1. What do you think about the weather today?<br>2. How often do you come to UCLA campus?<br>3. How did you celebrate last Halloween?<br>4. How often do you get your hair cut? Where do you go? Have you ever had a really bad haircut experience?<br>5. What is the best TV show you've seen in the last month? Tell your partner about it.<br>6. When was the last time you walked for more than an hour? Describe where you went and what you saw.<br>7. Do you like to get up early or stay up late? Why?<br>8. Do you have anything planned for later today? When are you going to do it?<br>9. Can you describe a conversation you had with another person earlier today?<br>10. What's your daily routine like?<br>11. What's the most stupid thing you have seen on TikTok?<br>12. What's your favorite exercise?<br>13. What's the last movie you saw in the theater?<br>14. What's your favorite social media?<br>15. What's your favorite meme?<br>16. What are your three favorite songs at the moment?<br>17. What restaurants do you recommend for friends from out of town?<br>18. Do you like hats? What kind?<br>19. What's your sign in astrology?<br>20. Do you prefer the beach or the mountain?<br>21. You have answered all the questions the experimenter provides. Feel free to talk about anything you want until the time is up! |
| Deep Condition | 1. What would constitute a "perfect" day for you?<br>2. Where is somewhere you've visited that you felt really had an impact on who you are today?<br>3. If you were going to become a close friend with the other participant, please share what would be important for him or her to know.<br>4. If a crystal ball could tell you the truth about yourself, your life, the future, or anything else, what would you want to know?<br>5. For what in your life do you feel most grateful? Tell the other participant about it.<br>6. Is there something you've dreamed of doing for a long time? Why haven't you done it?<br>7. What is one of the more embarrassing moments in your life?<br>8. What is one of your most meaningful memories? Why is it meaningful for you?<br>9. Can you describe a time you cried in front of another person?<br>10. If you could undo one mistake you have made in your life, what would it be and why would you undo it?<br>11. You have answered all the questions the experimenter provides. Feel free to talk about anything you want until the time is up! |

Note: Questions 1-10 from both the shallow and deep conditions have been adapted from Kardas, Kumar, & Epley (2021). Additional questions were added to the shallow condition for sufficient content in a 20-minute discussion.

### fNIRS data acquisition and preprocessing

Participants were scanned using a mobile fNIRS system (NIRSport2 by NIRx Medical Technologies, LLC, NY). The probe layout was comprised of 16 light sources and 16 detectors with a 3-cm average source-detector separation distance, which forms 42 channels (source-detector pairs) for partial-brain coverage across mentalizing (i.e., medial prefrontal cortex (mPFC) and temporo-parietal junction (TPJ)) and working memory regions (i.e., lateral prefrontal cortex (lPFC) and superior parietal lobule (SPL)) (Fig 2). The montage layout (Fig 2) was created in accordance with the 10-10 UI external positioning system to ensure consistency across head sizes. We measured participants’ head sizes and then fitted them with caps of appropriate sizes that affix the optodes to the scalp. Raw light intensity data was collected at a sampling rate of 5.09 Hz at wavelengths of 760 and 850 nm.

**Figure 2.**
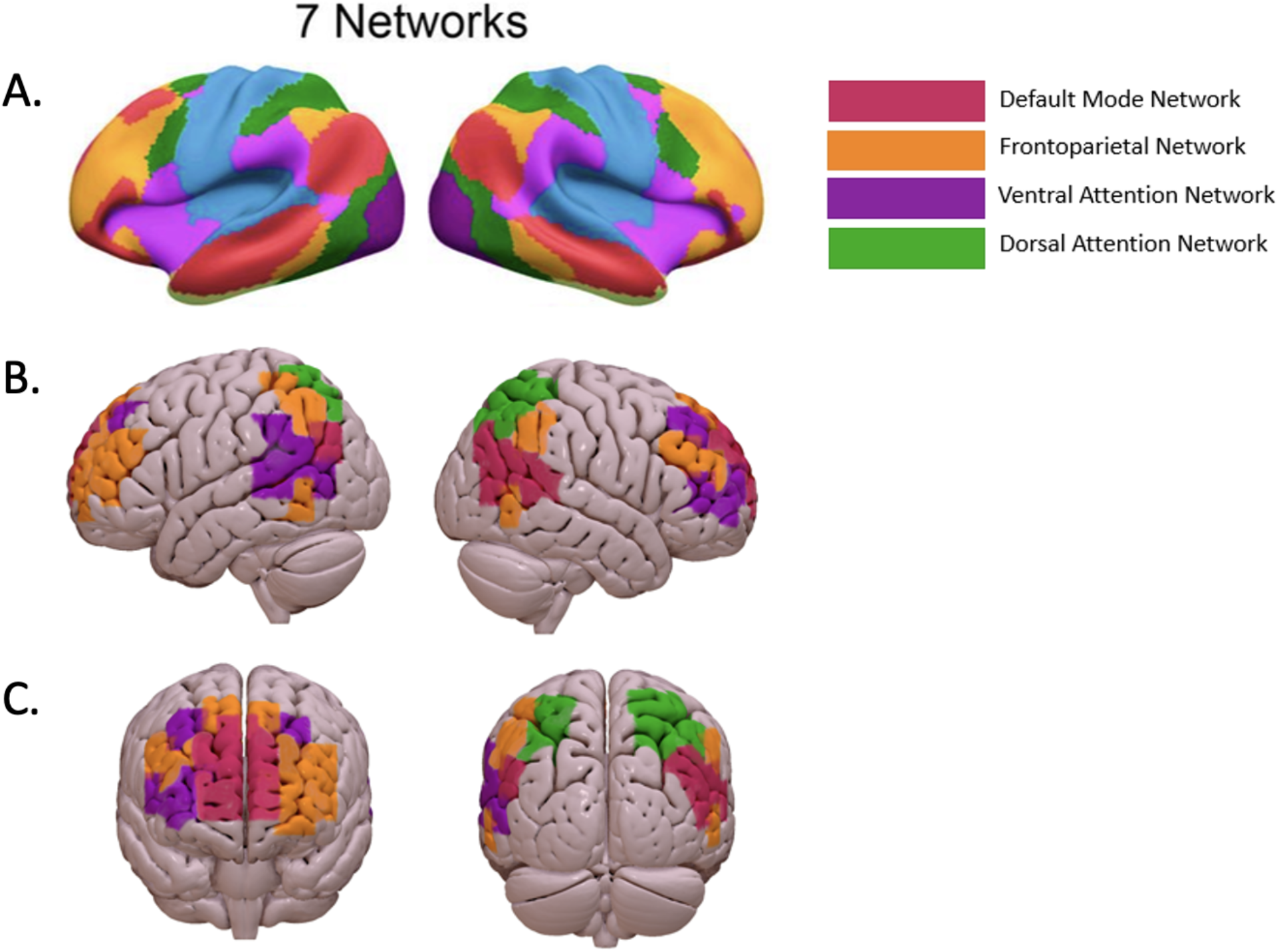
fNIRS montage consists of 40 channels for partial-brain coverage of cortical regions implicated in social interactions, including the default mode network (DMN) in red, the frontoparietal network (FPN) in orange, the ventral attention network (VAN) in purple, and the dorsal attention network (DAN) in green. A.Seven-network cortical parcellation defined by Yeo et al. (2011). B.Lateral views of DMN, FPN, VAN, and DAN. C.Anterior (left) and posterior (right) views of the same four networks.

Collected NIRS data underwent a comprehensive preprocessing pipeline. This pipeline utilized custom scripts in MATLAB alongside the Homer2 software suite (Huppert et al., 2009), adhering to established fNIRS best practices (Yücel, 2021). Emphasis was placed on analyzing oxyhemoglobin (HbO) concentrations, which prior research has indicated are more responsive to changes in cerebral blood flow than deoxyhemoglobin (HbR) levels work (Pan et al., 2017).

The preprocessing began with removing unrelated data – each time-course was truncated based on a trigger that indicated the start of the conversation. Noisy and oversaturated channels were identified and excluded using a modified quartile coefficient of dispersion (Bonett, 2006), with specific thresholds adjusted for the sampling rate (Cthresh = 0.6 – 0.03*sampling rate). Further refinement of the data included corrections for motion and non-neural changes in blood oxygenation. To address motion artifacts, discrete wavelet transform techniques (Molavi & Dumont, 2012) were performed to remove spike artifacts. To address non-neural physiological influences (e.g., cardiac and respiratory rhythms) and baseline drift, a conservative bandpass filter (0.008-0.2 Hz) was applied. Past work suggests that the cognitive dynamics of interest in this study are primarily manifested in lower frequency ranges (Sasai et al., 2011; Zuo et al., 2010).

Filtered data were then transformed from optical density to hemoglobin concentration values. This conversion used the modified Beer-Lambert Law (MBLL) with a standard differential path length filter [6, 6], commonly applied to adult cortical tissue to account for light dispersion. The final quality control step involved an autocorrelation change assessment to gauge the impact of motion correction. Channels displaying a substantial change in autocorrelation (exceeding a threshold of r = 0.1) were deemed significantly influenced by motion and thus excluded from subsequent analyses.

### Behavioral analyses for conversation and connection

Participants completed a post-conversation survey using a 5-point Likert scale, rating their experience in terms of bonding, engagement, liking, enjoyment, engagement, interest, awkwardness, ease, and perceived depth of the conversation; as well as epistemic knowledge, and similarity to the partner; and willingness for future interaction and the potential to become friends in real life.

We created the ‘connection’ composite variable by averaging the measures from three items (bonding, engagement, and liking) with greatest correlation between themselves (*r* > 0.6). We then calculated the mean within-dyad connection, which is used in both behavioral and neural analysis below.

To induce variability of connection across stranger dyads, we introduced shallow and deep topics of conversation as experimental conditions. However, with the naturalistic nature of conversations, we expected dyads to show variability in conversation depth not fully captured by condition. Thus, we report behavioral findings based on condition assignment as well as on self-reported felt depth of the conversation and use each as predictors of connection.

### Neural synchrony across brain networks and anatomical ROIs

Yeo et al. (2011) identified a 7-network parcellation of the human cerebral cortex based on 1,000 subjects. These networks converged and extended on networks previously described in resting-state literature, including the default networks (DMN), the frontoparietal network (FPN), the ventral attention network (VAN), and the dorsal attention network (DAN). We first divided the cortical areas covered by the 40 channels in our fNIRS montage (Fig. 2) based on the Yeo parcellation. We then obtained neural synchrony scores by computing the Pearson’s correlation for the average time series for each participant within each of the networks.

Since our main hypothesis focused on DMN, we also analyzed sub-regions within the DMN (namely, medial PFC, left TPJ, and right TPJ) to further understand the brain responses. For these analyses, we strictly separated lateralized ROIs and disregarded 2 channels on the border of medial PFC and lateral PFC.

### Predicting felt connection by neural synchrony in DMN

We used linear regression to examine the extent to which the four neural networks (DMN, FPN, VAN, DAN) predicted connection, and then examined which lateralized ROIs DMN predicted connection. We then integrated the neural findings with the behavioral findings related to felt depth by conducting a linear regression analysis with DMN neural synchrony and condition assignment, as well as DMN neural synchrony and felt depth as predictors of connection.

We also used a machine learning approach with binary classification (MATLAB’s fitclinear function, Version 2022b, The MathWorks, Inc.) to examine if high and low connection dyads could be predicted based on network synchrony and felt depth. All 70 dyads were categorized into high (N = 35) and low (N = 35) connection groups based on their average connection scores, with scores above 3.33 classified as high and those below 3.17 as low. A 5-fold cross-validation strategy was employed to mitigate overfitting, wherein the dataset was randomly partitioned into approximately five equal folds. For each iteration, four folds were used for training, which comprised 80% of the dataset, while the remaining fold, which was 20% of the dataset, served as the test set. To enhance the robustness of classification performance estimates, the entire cross-validation procedure was repeated 1,000 times with different random partitions. Model performance was quantified by mean classification accuracy, averaged across all repetitions and cross-validation folds, with standard deviation computed to assess variability. To determine if the model was significant the mean accuracy of the predictive model was compared to a null distribution which was created by randomly shuffling the outcome of connection for 1,000 repetitions.

### Data and Code Accessibility

The general fNIRS pre-processing pipeline can be accessed at https://github.com/abinnquist/fNIRSpreProcessing.

Data and code specific to this paper can be accessed at https://github.com/miaoqy0729/ConnectionStrangerDMN.

## Results

### Behavioral Results

The average self-reported connection for the full sample (*M* = 3.30) was significantly higher than the composite connection midpoint (*M* = 3.00) of “somewhat”, meaning dyads were closer to “very” connected than “not at all” connected (*t(104)* = 6.62, *p* < .0001). Replicating Kardas, Kumar, & Epley (2021), those in the deep conversation condition (*M* = 3.42) felt more connected than those in the shallow conversation condition (*M* = 3.18) (*t(103)* = −2.71, *p* = .008) (Fig 3. *left*). This same general trend held true for the 70 dyads whose fNIRS data was collected. Self-reported connection was significantly higher than the composite midpoint (*t(68)* = 5.44, *p* < .0001) and though marginal, those in the deep condition (*M* = 3.41) felt more connected than those in the shallow condition (*M* = 3.20) (*t(68)* = 1.96, *p* = .055). All additional analyses will focus on these 70 dyads.

**Figure 3.**
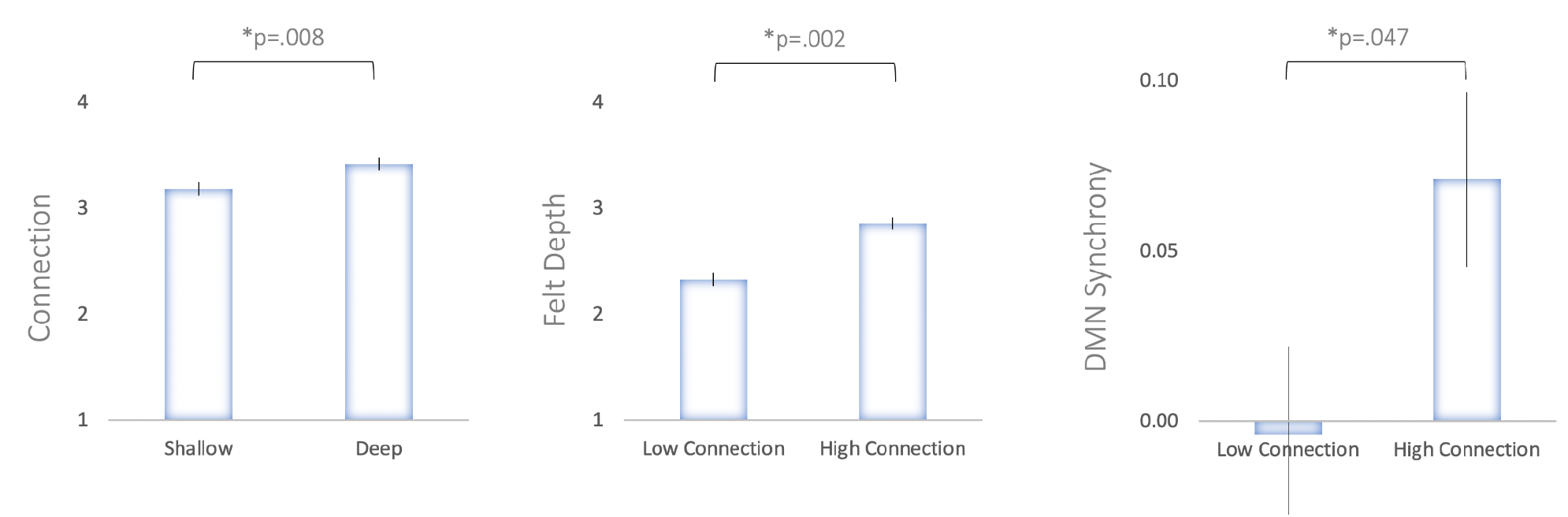
Difference across groups in behavioral and neural signals *Left*: Deep conversation dyads reported significantly greater felt connection after conversation than shallow conversation dyads. *Middle*: High connection dyads reported greater felt depth of the conversation than low connection dyads. *Right*: High connection dyads generated greater neural synchrony in the default mode network (DMN) during conversation than low connection dyads.

Although we assigned participants to shallow and deep conversation conditions, 28.6% of dyads (20 of 70 dyads) reported felt depth that was inconsistent with this assignment. Specifically, 8 of the 36 dyads in the shallow condition reported higher than average conversation depth and 12 of the 34 dyads in the deep condition reported lower than average conversation depth. Like the depth condition effect above, when mean-split into high vs. low felt depth groups, the same trend emerged: high self-reported depth was associated with more self-reported connection (*M* = 3.54) than low self-reported depth (*M* = 3.12) (*t*(*6*8) = −4.21, p < .001). Additionally, felt depth correlated strongly with self-reported feelings of connection after the conversation (*r*(70)=.55, p < 0.0001). Finally, mean-splitting on high vs. low self-reported connection produced analogous results: high connection dyads rated their conversations as significantly deeper (*M* = 2.86) than those low connection dyads (*M* = 2.33) (*t*(68) = –3.31, *p* = .002) (Fig 3. *middle*).

### Neural Results

We used linear regression to examine the relationship between dyadic neural synchrony in each of the four neural networks (DMN, FPN, VAT, DAT) and self-reported connection in dyads. As hypothesized *a priori*, DMN neural synchrony significantly predicted self-reported connection after the conversation (*β*= 0.874, *t(67)* = 2.51, *p* = .0146). Consistently, a Pearson correlation also revealed a positive association between DMN synchrony and connection (*r(68)* = 0.29, p = .015) (Fig 4. *left*). DMN neural synchrony also significantly predicted condition assignment as revealed by a logistic regression (*β*= −3.76, *z* = −2.15, *p* = .031). Participants who reported high connection also exhibited significantly greater DMN synchrony (*M* = 0.071) than those who reported low connection (*M* = −0.004) (*t*(67) = 2.03, p = .047) (Fig 3. *right*). In contrast, neural synchrony in other networks did not predict connection, including FPN (*β*= 0.546, *t(67)* = 1.40, *p* = .166), VAT (*β*= 0.614, *t(68)* = 1.52, *p* = .134), and DAT (*β*= 0.785, *t(47)* = 1.13, *p* = .264). Somewhat unexpectedly, DMN neural synchrony did not predict self-report felt depth after the conversation (*β*= −0.28, *t(67)* = −0.49, *p* = .623).

**Figure 4.**
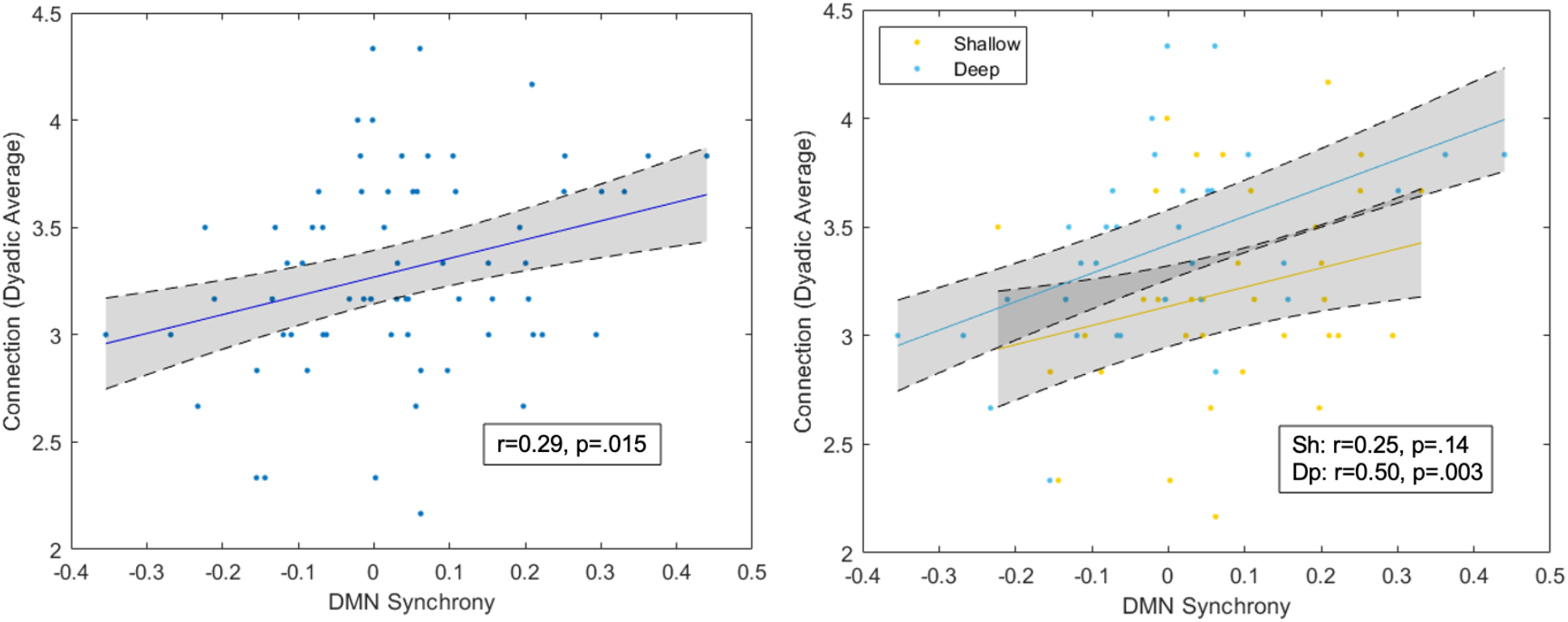
Correlations between neural synchrony in the default mode network (DMN) and key dependent variable connection *Left*: DMN synchrony is positively correlated with self-reported connection. *Right*: The correlation between DMN synchrony and connection is stronger in the deep condition compared to the shallow condition.

To further understand the brain responses within the DMN, we conducted analyses on each of DMN subregions including medial prefrontal cortex (mPFC) and both left and right temporoparietal junction (TPJ). Neural synchrony in mPFC predicted self-reported connection (*β*= 0.900, *t(68)* = 2.42, *p* = .0182). Neural synchrony in right TPJ was the strongest predictor of self-reported connection (*β*= 1.485, *t(64)* = 3.12, *p* = .0027) of any neural metric we examined, whereas neural synchrony in left TPJ did not significantly predict self-reported connection (*β*= 0.759, *t(66)* = 1.66, *p* = .102).

A regression model including experimental condition (shallow/deep) and DMN synchrony significantly predicted dyadic connection (R^2^_adjusted_ = 0.158, *F(2,66)* = 7.39, *p* = .0013). Within this model, condition (*β*= 0.300, *t(66)* = 2.80, *p* = .0067) and DMN synchrony (*β*= 1.134, *t(66)* = 3.29, *p* = .0016) both explained a significant amount of variance for predicting connection. In the smaller subsamples of the shallow (n = 36) and deep (n = 34) dyads, we observed DMN synchrony correlated with felt connection in the deep condition (r = 0.50, p = .003), but not in the shallow condition (r = 0.25, p = .14) (Fig 4. *right*). A correlation between connection and synchrony was also observed for the deep condition in FPN (r = 0.38, p = .029) and VAT (r = 0.35, p = .044) while neither was significantly correlated with connection in the shallow condition for FPN (r = 0.06, p = .745) and VAT (r = 0.11, p = .527).

Given that a number of dyads reported feelings of depth that were out of alignment with their experimental condition, we conducted an additional regression model that examined self-reported felt depth and DMN as predictors of connection regardless of condition. Results showed that the model significantly explains variance in connection (R^2^_adjusted_ = 0.368, *F(2,66)* = 20.8, *p* < .0001). Within this model, felt depth (*β*= 0.357, *t(66)* = 5.69, *p* < .0001) and DMN synchrony (*β*= 0.973, *t(66)* = 3.38, *p* = .0012) both explained a significant amount of variance for predicting connection.

### Classification Results

We next conducted a machine learning classification analysis using logistic regression with 5-fold cross-validation in order to classify dyads into low and high self-reported connection. Firstly, DMN neural synchrony and condition assignment were used as features to predict self-reported connection. We observed accurate classifications of 60.2% across 1,000 repetitions (*p* = .07). Then, DMN neural synchrony and self-reported depth were used as features to predict self-reported connection. We observed accurate classifications of 64.5% across 1,000 repetitions (*p* = 0.012). When DMN neural synchrony alone was used as the feature to predict self-report connection, we observed accurate classifications of 55.0% across 1,000 repetitions (*p* = .19).

Using neural synchrony in mPFC, left TPJ, and right TPJ as features to predict self-report connection, we observed accurate classifications of 61.4% across 1,000 repetitions (p = .04). When mPFC neural synchrony alone was used as the feature to predict self-report connection, we observed accurate classifications of 53.3% across 1,000 repetitions (*p* = .36). When left TPJ neural synchrony alone was used as the feature to predict self-report connection, we observed accurate classifications of 48.6% across 1,000 repetitions (*p* = .54). When right TPJ neural synchrony alone was used as the feature to predict self-report connection, we observed accurate classifications of 62.6% across 1,000 repetitions (*p* = .01).

## Discussion

Social neuroscience has made great strides in understanding how our social nature is instantiated in patterns of brain activity. Yet much of this work has focused on individuals in isolation—lying in MRI scanners, observing social stimuli, or imagining interpersonal experiences from a distance. With the growing adoption of more ecologically valid methods, relying on techniques like functional near-infrared spectroscopy (fNIRS), we can now examine real-time brain activity during face-to-face social interactions, capturing the dynamic and reciprocal nature of human connection.

In this study, we conducted the first investigation of how neural synchrony during a get-to-know-you conversation between strangers relates to the subjective feeling of connection after the conversation. We hypothesized that synchrony in the default mode network (DMN) would predict post-conversation feelings of connection. Across multiple analyses, including machine learning classification, this hypothesis was supported. At a more granular level, synchrony within the right TPJ subregion of the DMN was the best predictor and classifier of feelings of connection. Additionally, both experimentally manipulated and self-reported measures of conversation depth were significantly related to connection, replicating past work (Kardas, Kumar, & Epley, 2021).

Why does DMN synchrony predict the formation of social connection? Growing evidence suggests that DMN synchrony, especially in TPJ, is a signal indicating that two or more people are subjective construing things in more similar ways (Lieberman, 2022; Lieberman, in press). There is already abundant evidence from shared reality theory suggesting that shared interpretations and seeing eye-to-eye make connection formation more likely to occur (Pinel et al., 2006; Launay & Dunbar, 2015; Rossignac-Milon et al., 2021). Prior to this study, it was already known that people with existing strong social connections are more likely to show DMN synchrony when watching video clips (Parkinson, Kleinbaum, & Wheatley, 2018; Li et al., 2022). However, this had not been examined in hyperscanning settings as strangers form (or do not form) connections through conversation. This study represents the first evidence that DMN synchrony operates in an analogous way when strangers have a get-to-know-you conversation and that DMN synchrony may be useful as an indicator of connection formation.

At first glance, these results may seem to be at odds with the recent results from Speer et al. (2024). They found that friends showed greater divergence in neural state space than strangers as they spoke to one another in fMRI scanners. While neural synchrony and movement through neural state space are distinct measures, they are not orthogonal; perfect synchrony throughout the brain would necessitate convergence in neural state space as well. It is important to note that the differences between strangers and friends in their study may reflect pre-existing differences in these groups in addition to within task dynamics. It may also be the case that the dynamics that predict successful conversations among friends may be different than those corresponding to strangers coming to feel more connected. Additionally, Speer et al. found that even among strangers, those who diverged more in neural state space *enjoyed* the conversation more, but this apparently did not translate to greater feelings of *closeness*, which would be most analogous to our measure of connection.

As with most initial findings, ours raise as many questions as they answer suggesting the need for additional research. For instance, we expected DMN neural synchrony to correlate with the self-reported depth of the conversation, but it did not. This raises the possibility that DMN neural synchrony might come in qualitatively different varieties, only some of which might be valid indicators of connection formation. For instance, there might be a difference between ‘easy’ and ‘hard’ DMN synchrony. Easy synchrony might occur during shallow conversations when discussing simple events for which both members of the dyad have strong familiarity. If a student describes walking along a path on campus to another student from the same campus, they can easily follow along despite this having little relevance for connection formation. To this end, in our results, DMN synchrony during the shallow condition did not significantly correlate with felt connection.

In contrast, hard synchrony would involve effortful co-construction of meaning during emotionally or cognitively complex conversations. Sharing a painful memory or revealing a personal truth requires more engagement and mutual attunement to reach alignment. DMN synchrony here may be a stronger signal of connection formation and in our results we saw a robust correlation between DMN synchrony and felt connection. While intersubject correlation may not be able to distinguish these two varieties of DMN synchrony, it is possible that other methods like wavelet transform, cross recurrence quantification analysis (cRQA; Coco & Dale, 2014), or a qualitative examination of quantitative time series (Miao et al., 2025) can.

A second open question is whether synchrony methods may be able to better pinpoint the precise moments that the connection between two people is growing or fading. If neuroimaging methods can be used to identify these moments, we can then try to reverse engineer what is being said or the nonverbal behavior that accompanies the words to improve our general understanding of the processes that facilitate or inhibit connection formation. Countless articles and blog posts make pronouncements about the best ways to make friends, but little is known about the key moments in first conversations that are critical to moving people towards or away from a closer connection. A sliding window approach to neural synchrony might be helpful, however, non-linear methods designed to extract dynamic structure, such as cRQA, might be promising here as well.

More broadly, the current work may inform intervention strategies designed to promote deeper connection in everyday life. Prior studies suggest that people often mispredict the outcomes of talking to strangers, underestimating how enjoyable or meaningful these conversations can be, and overestimating the likelihood of awkwardness (Boothby et al., 2018; Kardas, Kumar, & Epley, 2021). Our findings lay the groundwork for potential interventions— such as brief educational prompts about the liking gap, misprediction, or importance of audacity—could encourage people to initiate deeper conversations by lowering social barriers and fostering conditions for more meaningful DMN synchrony.

Such work could also examine other key factors like participant personality and current level of loneliness. There are also open questions about the extent to which DMN is a critical marker of seeing eye-to-eye in other contexts such as cross-ideological conversations, negotiations, classroom education, therapy, performance within teams, and even interactions with AI agents. Future work is needed to determine the breadth and limits of application of fNIRS-based DMN synchrony during social interactions.

## Conclusion

This study provides the first direct evidence that neural synchrony within the default mode network (DMN), particularly in the right TPJ, predicts the formation of interpersonal connection during naturalistic conversations between strangers. Using fNIRS hyperscanning and machine learning, we demonstrated that DMN synchrony, in conjunction with perceived conversational depth, reliably distinguished high-from low-connection dyads. These findings extend prior work on DMN synchrony during passive co-viewing to real-time, face-to-face interactions, supporting the hypothesis that DMN synchrony functions as a neural index of “seeing eye-to-eye.”

More broadly, our results underscore the value of deep conversations in promoting social bonding and offer a mechanistic account of how such bonds may arise through shared neural representations. As loneliness becomes an increasingly pressing public health concern, these findings suggest novel avenues for fostering connection through structured interactions and neural feedback. Future work should explore the temporal dynamics of synchrony during dialogue and test the generalizability of these effects across more diverse populations and interaction contexts.

